# The impact of plasma membrane lipid composition on flagella-mediated adhesion of enterohemorrhagic *Escherichia coli*

**DOI:** 10.1101/2020.07.07.189852

**Authors:** Hélène Cazzola, Laurine Lemaire, Sébastien Acket, Elise Prost, Luminita Duma, Marc Erhardt, Petra Čechová, Patrick Trouillas, Fady Mohareb, Claire Rossi, Yannick Rossez

## Abstract

Enterohemorrhagic *Escherichia coli* (EHEC) O157:H7 is a major cause of foodborne gastrointestinal illness. The adhesion of EHEC on host tissues is the first step enabling bacterial colonization. Adhesins like fimbriae and flagella mediate this mechanism. Here, we studied the interaction of the bacterial flagellum with the host cell’s plasma membrane using Giant Unilamellar Vesicles (GUVs) as a biologically relevant model. Cultured cell lines contain many different molecular components including proteins and glycoproteins. In contrast, with GUVs we can characterize the bacterial mode of interaction solely with a defined lipid part of the cell membrane. Bacterial adhesion on GUVs was dependent on the presence of the flagellar filament and its motility. By testing different phospholipid head groups, the nature of the fatty acid chains or the liposome curvature, we found that lipid packing is a key parameter to enable bacterial adhesion. Using HT-29 cells grown in the presence of polyunsaturated fatty acid (α-linolenic acid) or saturated fatty acid (palmitic acid), we found that α-linolenic acid reduced adhesion of wild type EHEC but not of a non-flagellated mutant. Finally, our results reveal that the presence of flagella is advantageous for the bacteria to bind to lipid rafts. We speculate that polyunsaturated fatty acids prevent flagellar adhesion on membrane bilayers and play a clear role for optimal host colonization. Flagella-mediated adhesion to plasma membranes has broad implications to host-pathogen interactions.

**Importance:** Bacterial adhesion is a crucial step to allow bacteria to colonize their hosts, invade tissues and form biofilm. Enterohemorrhagic *E. coli* O157:H7 is a human pathogen and the causative agent of diarrhea and hemorrhagic colitis. Here, we use biomimetic membrane models and cell lines to decipher the impact of lipid content of the plasma membrane on enterohemorrhagic E. *coli* flagella-mediated adhesion. Our findings provide evidence that polyunsaturated fatty acid (α-linolenic acid) inhibits E. *coli* flagella adhesion to the plasma membrane in a mechanism separate from its antimicrobial and anti-inflammatory functions. In addition, we confirm that cholesterol-enriched lipid microdomains, often called lipid rafts are important in bacterial adhesion. These findings significantly strengthen plasma membrane adhesion via bacterial flagella in an important human pathogen. This mechanism represents a promising target for the development of novel anti-adhesion therapies.

## Introduction

Enterohemorrhagic *Escherichia coli* (EHEC) serotype O157:H7 are Shiga-toxin producing strains characterized by peritrichous flagella and is responsible for major food-borne diseases and for serious infections (1). When ingested, infection with EHEC is characterized by symptoms ranging from hemorrhagic colitis to life-threatening complications (2). These bacteria have the capacity to infect and to multiply in a wide variety of host species including human, animals and even plants (3). The persistence in their hosts, including humans, occurs through adhesion onto tissues (4). EHEC adhere to the intestinal mucosa in a manner termed the attaching and effacing effect (5) but other mechanisms, involving flagella and pili, have been described but not fully characterized (6). The pili are the most described adhesins present at the bacterial surface (7, 8). More recently, bacterial flagella have also been identified in bacterial adhesion on different host tissues (9, 10). The bacterial flagellum is a multiprotein complex, best known as a filament responsible for bacterial movement toward preferred environmental niches (11). The presence of flagella can be seen as a characteristic marker of early-stage colonization.

The flagellum is mainly composed of a globular protein, the flagellin, which is organized in four connected domains named D0, D1, D2 and D3. Flagellin peptides fold back on themselves and the D0-D1 domains are interacting through a coiled-coil interface and hydrophobic contacts, which are essential in the flagellin polymerization process. These N- and C-terminal regions are well conserved across all bacterial flagellins. Conversely, the D2-D3 domains generate antigenic diversity and are exposed on the filament exterior (12, 13). These monomers form a helix made of 11 protofilaments of flagellin (14) with lengths up to 15-20 μm and a diameter around 20 nm.

Recent evidence has suggested that flagella bind to the plasma membrane phospholipids mainly through hydrophobic effects (15, 16). However, little is known about the parameters that govern the direct interaction between the lipid bilayers of plasma membranes and the flagella of enteropathogenic bacteria. Previously, membrane rafts, which are membrane domains enriched in cholesterol and sphingolipids (17, 18), have been documented as targets of bacterial pathogens when targeting host membranes (19–22). To shed light on the impact of lipid composition and physical properties of the lipid bilayer on the adhesion of O157:H7 flagella onto the host cell’s plasma membranes, we selected a series of biomimetic membranes to study the impact of phospholipid polar head group, lipid size or fatty acid saturation. Various membrane parameters were tested including, fluidity, area per lipids, membrane thickness and order parameters by biophysical techniques and molecular dynamics (MD) simulations. Here, we demonstrate that bacterial flagella from EHEC O157:H7 can bind to the cellular plasma membrane by exploiting membrane fluidity and head group packing. We further found that polyunsaturated fatty acids reduce flagellar adhesion and lipid rafts promote flagellar adhesion.

## Results

### Interactions of bacterial flagella with phosphatidylcholine vesicles

To expand our knowledge about bacterial flagellum adhesion on plasma membranes (15, 16), we initiated this study by visualizing the interaction of bacteria with biomimetic GUVs composed mainly of phosphatidylcholine (PC) from eggs (egg-PC) at room temperature (23°C). This was performed by combining fluorescence and phase contrast microscopy. The lipid bilayer was doped with 2% molar of a green-emitting fluorescent phospholipid, namely 1,2-dipalmitoyl-sn-glycero-3-phosphoethanolamine-N-(7-nitro-2-1,3-benzoxadiazol-4-yl) (NBD-PE) (Fig. 1A and 1B). The GUV average diameter was 6.27 μm with diameters ranging from 2 to 41 μm (calculated from more than 450 GUVs) (Fig. S1). Such sizes are relevant with that of EHEC host cells (5-30 μm in diameter). Following incubation with both non-flagellated and flagellated EHEC, the former bacteria were not easily found around the GUVs (Fig. 1A) whereas the latter were mainly observed around the GUVs (Fig. 1B).

**FIG 1.**
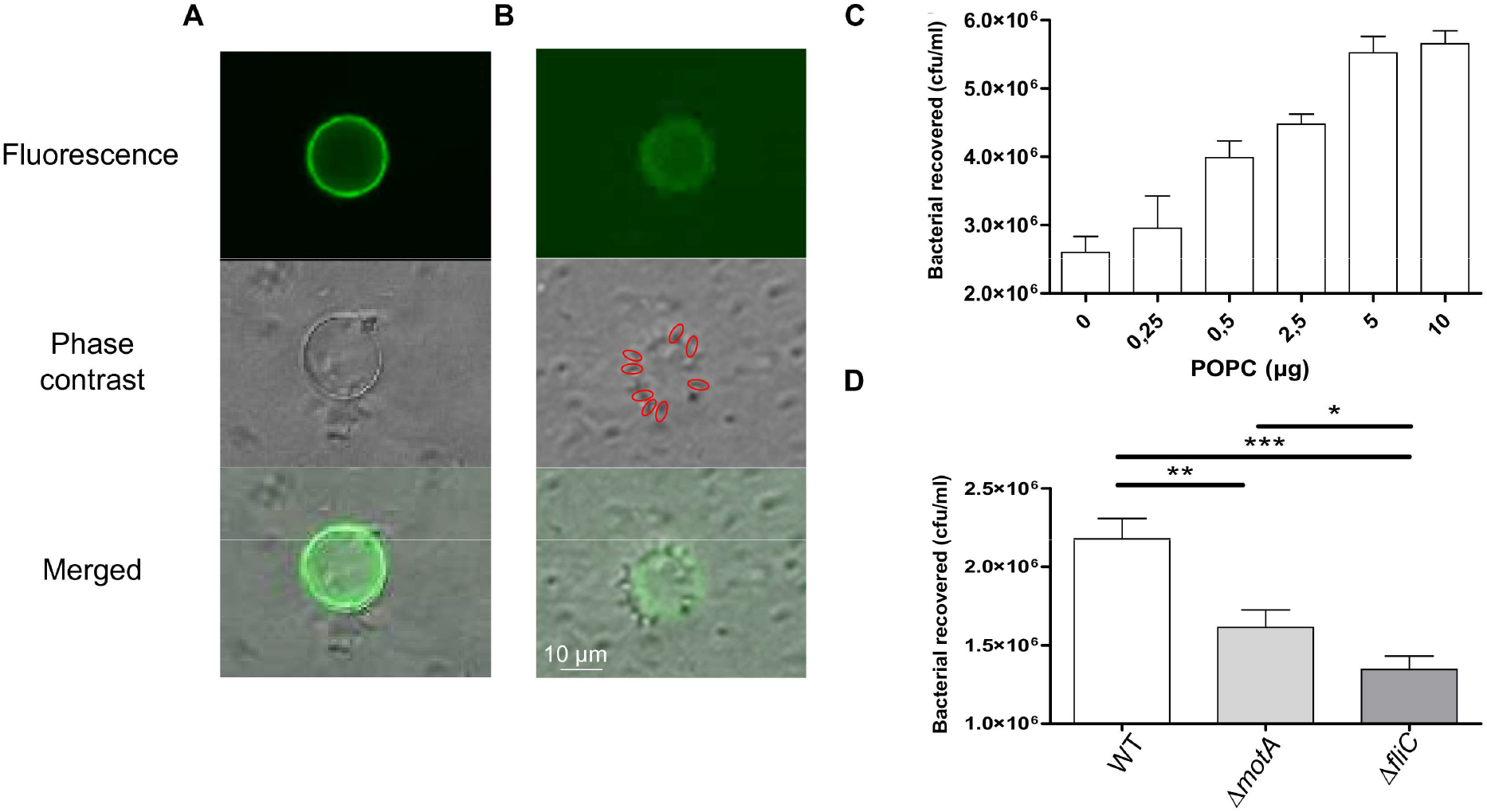
Interaction of egg-PC (GUV) with EHEC O157:H7. View of the liposome surface after immobilization (doped with 2% molar NBD-PE) (A) EHEC Δ*fliC* and (B) EHEC WT. Bacteria are visible in phase contrast. Red circles show bacteria around the GUV. (C) Interaction of the same bacterial concentration on different lipid vesicle quantities (0, 0.25, 0.5, 2.5, 5 and 10 μg). n=9 biologically independent samples. (D) Adhesion of EHEC WT, Δ*motA* and Δ*fliC.* Non-specific adhesion on plastic was subtracted. WT: n=51; ΔmotA: n=31; Δ*fliC:* n=18 biologically independent samples. The bar graphs represent the mean of the reported data. Statistical significances were determined by a two-tailed Student’s *t* test and are defined as follows: ***: *p* ≤ 0.001; **: *p* ≤ 0.005; *: *p* ≤ 0.05. The bar graphs represent the mean of the reported data.

The Brownian motion of GUVs and the motility of bacteria make the interaction between the two species difficult to quantify by microscopy. Therefore, we used a quantitative binding assay on GUVs as described previously (16) and illustrated in Fig. S2.

The lipid quantity immobilized on the GUVs’ surface was dose-dependent (Fig. 1C). Lipid GUV suspension containing from 0 to 10 μg of PC were deposited onto the gold surface and subsequently incubated with EHEC. An increase of adherent bacteria was observed up to 5 μg. This plateau means that saturation of bacterial adhesion was achieved under the experimental conditions. As a result, all the other experiments were performed with 5 μg of GUVs.

Both the presence of the flagella and their motility was assessed to understand the role of flagella in bacterial adhesion. The comparison between EHEC wild type (WT) and the flagella-free EHEC mutant *(ΔfliC)* confirmed that the lack of flagella is responsible for less adherent bacteria (Fig. 1D). A non-motile but flagellated EHEC mutant *(ΔmotA)* was also studied. EHEC Δ*motA* was more adherent than Δ*fliC* (Fig. 1D), suggesting that the flagellar movement is essential to bacterial adhesion. Collectively, these results confirmed the suitability of the GUV adhesion assay to investigate the adhesive properties of EHEC flagella and reveal a key role of bacterial flagella and active motility in the adhesion process on plasma membrane lipid bilayers.

### The role of vesicle curvature in flagellar adhesion

To tackle the impact of curvature on flagellar adhesion, Large Unilamellar Vesicles (LUVs) were produced with a diameter of 400 ± 30 nm. Surprisingly, EHEC were less adherent on LUVs than on GUVs (Fig. 2A), suggesting an impact of the size of the vesicles, thus membrane curvature, on flagellar adhesion. To further investigate vesicle curvature-dependence, generalized polarization (GP) measurements were achieved with Laurdan (6-dodecanoyl-2-dimethylaminonaphthalene) an amphiphilic fluorescent dye sensitive to local packing. Because Laurdan GP sensed changes in the phospholipid order (23, 24), if the bacterial flagella can penetrate the plasma membrane, a lower GP order will be observed (Fig. 2B). Vesicles with different sizes were incubated with purified H7 flagella to investigate the impact of four different bilayer curvatures. The GP measurement could not be performed on the whole bacteria because the probe detects mainly the bacterial motility in the surrounding environment of the vesicles (data not shown). As the initial packing of the lipid bilayers depends on the curvature, the initial GP value of the vesicles with different diameters were normalized by subtracting the GP value in the absence of flagella for each size of liposome (ΔGP) (Fig. 2C). The presence of H7 with vesicles allowed slight changes in the lipid order only for vesicle diameters greater than 2 μm. This confirms the existence of a threshold diameter above which the interaction becomes significant.

**FIG 2.**
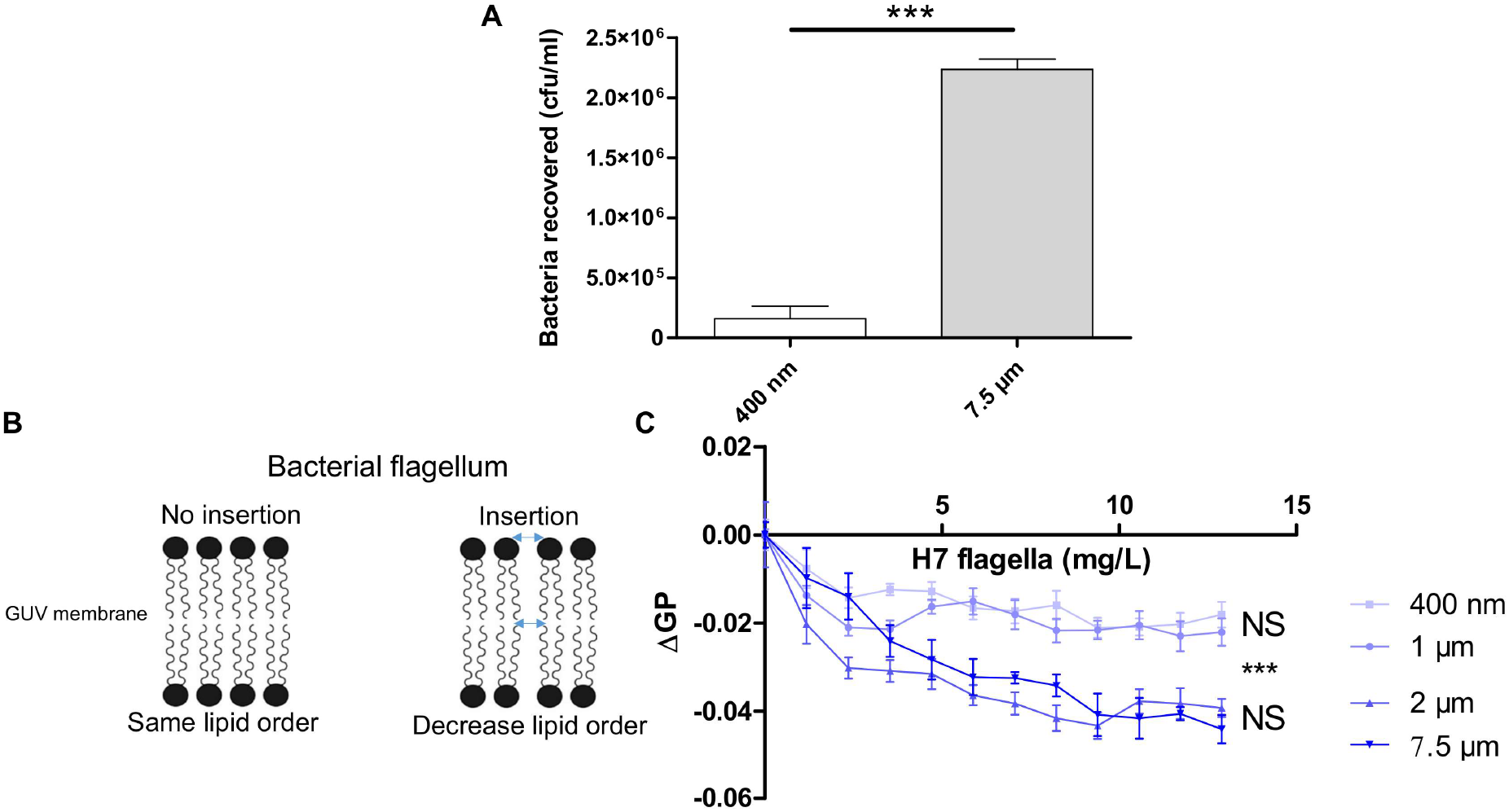
Bacterial flagella adhesion and lipid vesicle curvature. (A) EHEC WT adhesion on phosphatidylcholine (egg-PC) GUVs (7.5 μm) and LUVs (~400 nm diameter). Non-specific adhesion on plastic was subtracted. 400nm: n=18; 7.5μm: n=51 biologically independent samples. (B) Scheme of the principle of the Generalized Polarization (GP) measurements. (C) Relative Laurdan emission ΔGP was calculated by successive addition of purified H7 flagella in solution containing different sizes of phosphatidylcholine (egg-PC) liposomes at 37 °C. For each concentration at least n=5 biologically independent samples were done. Statistical significances were determined by a two-tailed Student’s *t* test and were calculated at the highest concentration of H7 flagella. The bar graphs represent the mean of the reported data. Significance levels are defined as follows: ***: *p* ≤ 0.001; NS: non-significant. The bar graphs represent the mean of the reported data.

### The role of phospholipid head groups in flagellar adhesion

We next evaluated more thoroughly how different lipid species may affect EHEC flagellar adhesion. Focus was first given to the influence of the lipid head groups as they could modulate bilayer curvature and lipid packing (25). Since our first results were obtained with vesicles composed of egg-PC, we characterized precisely the fatty acid profile and the lipid species of the egg-PC that we used by gas chromatography-flame ionization detection (GC-FID) and liquid chromatography-mass spectrometry (LC-MS) respectively. The main fatty acids were C16:0 (palmitic acid), C18:0 (stearic acid), C18:1 (oleic acid) and C18:2 (linoleic acid). PC 34:1, constituted of palmitic acid and oleic acid, named 1-palmitoyl-2-oleoylphosphatidylcholine or POPC was the most abundant lipid (Fig. S3).

In the following experiments, membranes made of POPC alone were compared to membranes made of POPC associated to other two phospholipids found in host plasma membrane with the same fatty acid composition: 1-palmitoyl-2-oleoyl-sn-glycero-3-phosphoethanolamine (POPE) and 1-palmitoyl-2-oleoyl-sn-glycero-3-phospho-(1’-rac-glycerol) (POPG). To prevent from destabilization of the bilayer, POPG and POPE were mixed with POPC at 60% mol (26). This allowed tackling the influence of the lipid head groups, which is known to impact on various bilayer’s properties including curvature or lipid packing (27). As POPC, POPE is zwitterionic but has a negative curvature. Conversely, POPG carries a negative charge at physiological pH and exhibits a zero spontaneous curvature as POPC (28).

A clear reduction of adhering bacteria was observed when POPC was mixed with POPE or POPG (Fig. 3A). To unravel the role of membrane fluidity, membrane thicknesses and area per lipids were calculated from MD simulations. The presence of POPE dramatically affected these membrane properties by decreasing the area per lipid and increasing membrane thickness. In other words, POPE increased ordering, as also seen by the order parameters of lipids in the POPE/POPC binary mixture with respect to the pure POPC bilayer (Fig. 3B). Conversely, the presence of POPG only slightly affected the area per lipid and increased membrane thickness. To further investigate membrane ordering, steady-state fluorescence anisotropy measurements were performed on GUVs (Fig. 3B) using two probes, diphenylhexatriene (DPH) and Laurdan (28). Their partitioning allows monitoring lipid dynamics in different membrane regions. The amphiphilic structure of Laurdan is localized at the hydrophobic-hydrophilic interface region whereas DPH locates in the hydrophobic core of the lipid bilayer (29).

**FIG 3.**
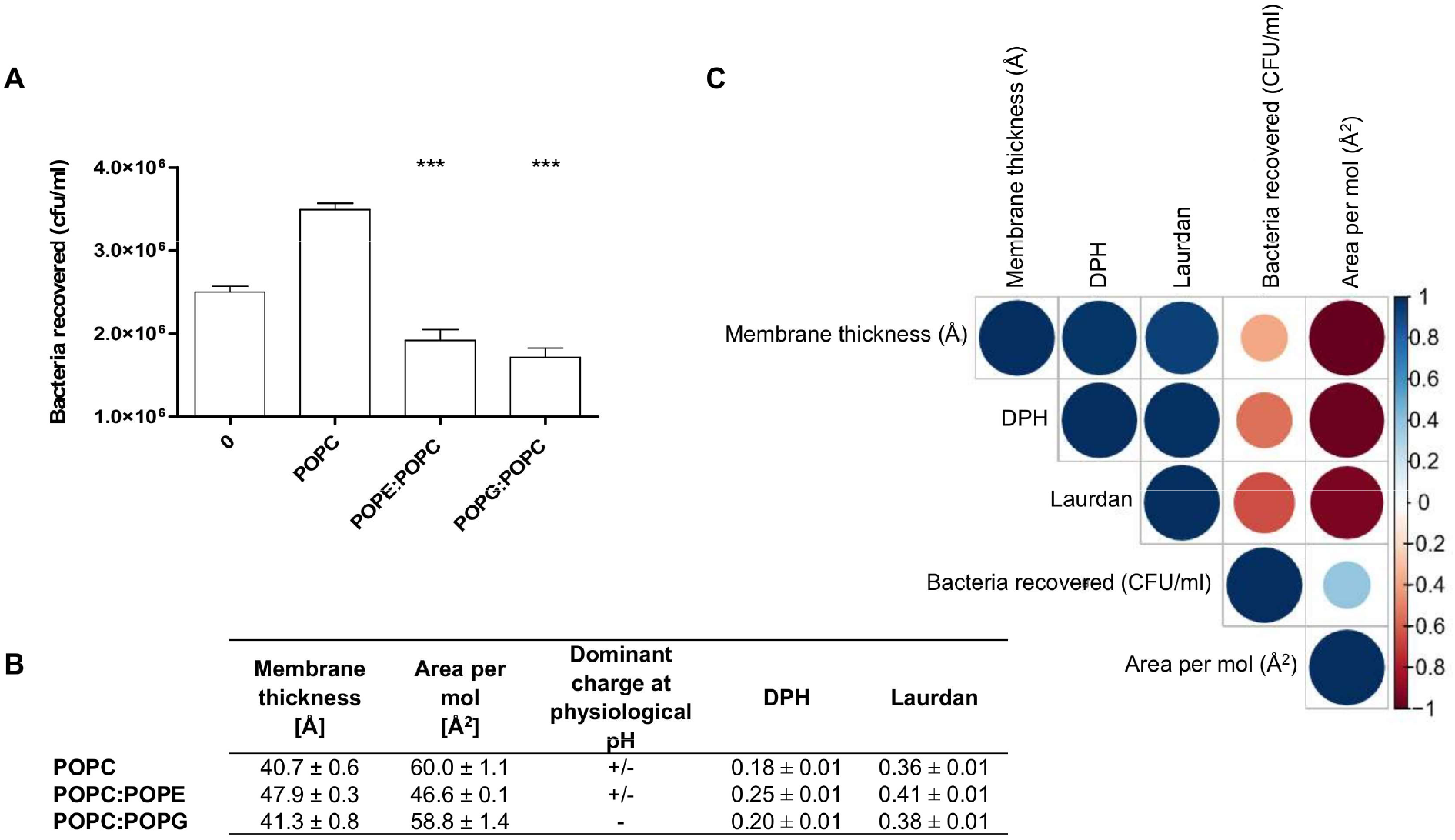
Impact of head polar phospholipids with one palmitic and one oleic acid (PO) on EHEC flagella adhesion. (A) EHEC adhesion on PO acyl chain lipids. 0 corresponds to the naked gold-coated glass surface. POPC:POPE and POPC:POPG were mixed 40:60 mol. Non-specific adhesion on plastic was subtracted. 0: n=135; POPC: n=25; POPC:POPE: n=12; POPC:POPG: n=23 biologically independent samples. The bar graphs represent the mean of the reported data. Statistical significances were determined by a two-tailed Student’s *t* test. Significance levels were defined as follows: ***: *p* ≤ 0.001. (B) Table summarizing anisotropy values for diphenylhexatriene (DPH) and Laurdan probes at 23 °C. The averages and standard deviations were calculated from at least three independent measurements. Membrane thickness and area per molecule, as well as related standard deviations, were calculated over the last 200 ns of MD simulations. (C) Correlogram representing Pearson’s correlation coefficients calculated for bacteria recovered, membrane thickness, area per mol, diphenylhexatriene (DPH) and Laurdan anisotropies for the following lipids: POPC, POPC:POPE, POPC:POPG. Positive correlations are visualized in blue while negative correlations are in red. The color intensity and the size are proportional to the correlation coefficients. In the right side of the correlogram, the legend color shows the correlation coefficients and the corresponding colors. More detailed information on the significance of the correlations as well as on the correlation coefficients can be found in Fig. S4.

As expected and already described in the literature (30), membrane thickness and area per lipid are negatively correlated. In addition, these parameters correlated with the fluidity values (Fig. 3C) showed a very high positive correlation between the two-bilayer order parameters. Furthermore, bacterial adhesion decreased when fluidity increased with a correlation coefficient ranging from −0.54 (for DPH values) to −0.65 (for Laurdan values). This correlation was clearly evidenced since POPC and POPE exhibit strong differences in their membrane properties, thickness and area per lipid (Fig. 3B). Conversely, the differences between POPC and POPG are much lower, however they are differently charged.

To further study the impact of the polar head group, a series of lipid bilayers made of two *cis*-oleic acid moieties, namely DOPC, DOPE and DOPG, was selected. Due to the presence of one unsaturation per lipid chain, the lipid order was significantly decreased compared to the PO-series, *i.e.,* greater membrane area per lipids and lower order parameters (Fig. 4A). As a consequence of this global disordering effect, the variations in the lipid order was smoothed within the DOPC, DOPE and DOPG series with respect to the PO-series, allowing a better focus on the sheer impact of the chemical feature and charge of the head groups. A synthetic positively charged lipid was also added: 1,2-dioleoyl-3-trimethylammonium-propane (DOTAP). The three lipids, DOPE, DOPG and DOTAP, were mixed with POPC at 40:60 molar ratio. Less bacteria were recovered on DOPC (~2.5 × 10^6^ CFU/ml) alone than with POPC (~3.0 × 10^6^ CFU/ml) (Fig. 4B). When mixing POPC with DOPC or DOPE, this number was significantly lower than DOPC alone. Interestingly, we observed no significant differences neither with DOPG nor with DOTAP mixed with POPC with respect to pure DOPC (Fig. 4B). The bacterial adhesion has a low negative correlation except for the area per lipid parameter, which is positive (Fig. 4C). An impact of the head group charge was observed on EHEC flagellum adhesion. Zwitterionic phospholipids for the DO-bilayer series decreased bacterial adhesion and conversely negative and positive head group had no impact. We thus reasoned that lipid charge and membrane properties could differentially affect flagella adhesion on plasma membrane lipids driven in part by the acyl chain content.

**FIG 4.**
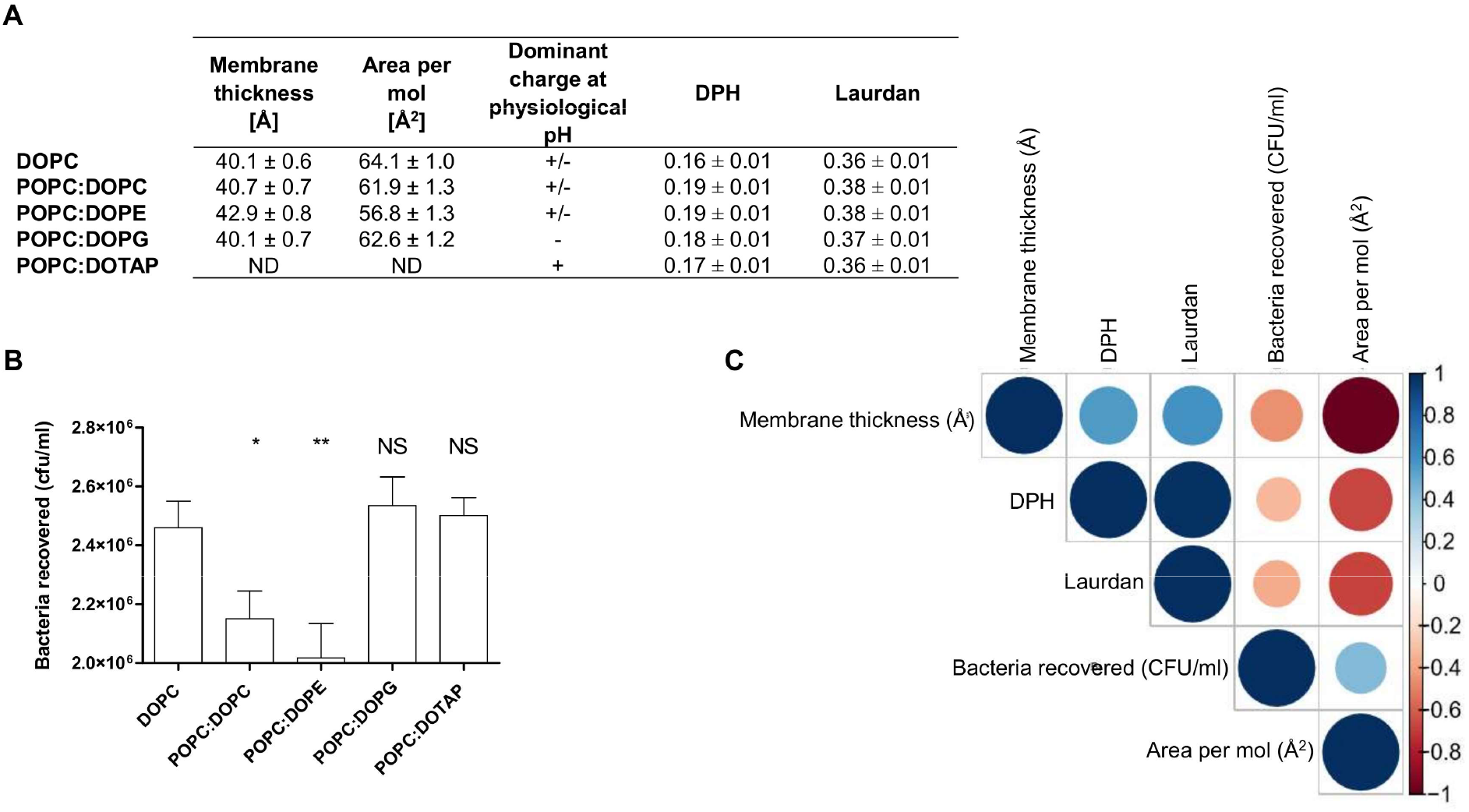
Impact of head polar phospholipids with two oleic acids (DO) on EHEC flagella adhesion. (A) EHEC adhesion on DO acyl chain lipids. All mixture compositions, except for DOPC, were POPC:X 40:60% mol with X the second lipid. Non-specific adhesion on plastic was subtracted. DOPC: n=13; POPC:DOPC: n=10; POPC:DOPE: n=19; POPC:DOPG: n=12; POPC:DOTAP: n=17 biologically independent samples. The bar graphs represent the mean of the reported data. Statistical significances were determined by a two-tailed Student’s *t* test. Significance levels are defined as follows: **: *p* ≤ 0.005; *: *p* ≤ 0.05; NS: non-significant. (B) Table summarizing anisotropy values for diphenylhexatriene (DPH) and Laurdan probes at 23 °C. The averages and standard deviations were calculated from at least three independent measurements. Membrane thickness and area per molecule, as well as related standard deviations, were calculated over the last 200 ns of MD simulations. (C) Correlogram representing Pearson’s correlation coefficient calculated for bacteria recovered, membrane thickness, area per mol, DHP and Laurdan anisotropies for the following lipids: DOPC, POPC:DOPC, POPC:DOPE, POPC:DOPG. Positive correlations are visualized in blue while negative correlations are in red. The color intensity and the size are proportional to the correlation coefficients. On the right-hand side of the correlogram, the legend color shows the correlation coefficients and the corresponding colors. More detailed information on the significance of the correlations as well as on the correlation coefficients can be found in Fig. S5.

### The role of phospholipid acyl chains in flagellar adhesion

To investigate further the impact of phospholipid unsaturation, directly related to the fluidity, on bacterial adhesion, the membrane parameters were assessed in the subsequent series: pure POPC, DOPC Δ9-cis (DOPC cis), DOPC Δ*9-trans* (DOPC *trans)* and a poly-unsaturated lipid, PC 18:3/18:3 (PC 18:3). All lipids were tested at room temperature as before (23 °C) because as long as we work above their phase transition temperature *(Tm),* no changes are introduced to the ordered gel phase. DOPC *trans* was also tested at 4 °C as it has a phase *Tm* of 12 °C. Namely at 23 °C, all lipids were in fluid (disordered) phase (Fig. 5A), whereas at 4 °C, the bilayer made of DOPC *trans* is ordered. At 23 °C, GUVs made of 100% DOPC exhibited less bacteria adhesion regardless of whether the double bond was in *trans* or *cis* compared to the POPC membrane (Fig. 5B). This is in line with their anisotropy under the same temperature condition. At 4 °C, DOPC *trans* is ordered and less bacteria can stick on it. Interestingly, the presence of two α-linolenic acids in the PC 18:3 drastically decreased adhesion, most probably because of a dramatic change in their fluidity and area per lipid, thus ordering. These results indicate that membrane fluidity can affect flagellar adhesion and extreme values (0.19 and 0.13 with DPH; and 0.33 and 0.45 with Laurdan) are not in favor of adhesion. The positive correlations between EHEC adhesion and the fluidity state of the bilayers for POPC, DOPC *cis,* DOPC *trans* and PC 18:3 at 23 °C were between high and very high (Fig. S6), indicating that the more fluid the membrane, the lower the bacterial interaction with the GUVs’ membrane. A correlation was performed to include all parameters (anisotropy values and values obtained by MD simulations) when available, in order to compare the same head group with different fatty acids (Fig. 5C). DOPC *cis,* POPC, POPC:DOPC and PC 18:2/18:3 at 23 °C were included. We must note here that bacterial adherence assays were obtained with PC 18:3 although MD simulation values were determined with PC 18:2/18:3. The former lipid is not available in simulation databases, whereas the latter lipid has already been parameterized in the Charmm force-field. These two lipids are structurally very close (same head group and only one differences in unsaturation) and very similar behavior of the lipid bilayer is likely (31). Moreover, anisotropy values were measured for PC 18: 2/18:2 (PC 18:2) and no difference was observed compare to PC 18:3 (Fig. 5C). All correlations between bacterial adhesion numbers and the other parameters are high except for Laurdan anisotropy that is moderate. To favor more adhering bacteria with pure PC GUVs, the membrane thickness needs to be around 40 Å and the area per molecule to 60 Å. A thinner plasma membrane and too much space between molecules do not help bacterial adhesion. These results are directly linked with those about membrane fluidity.

**FIG 5.**
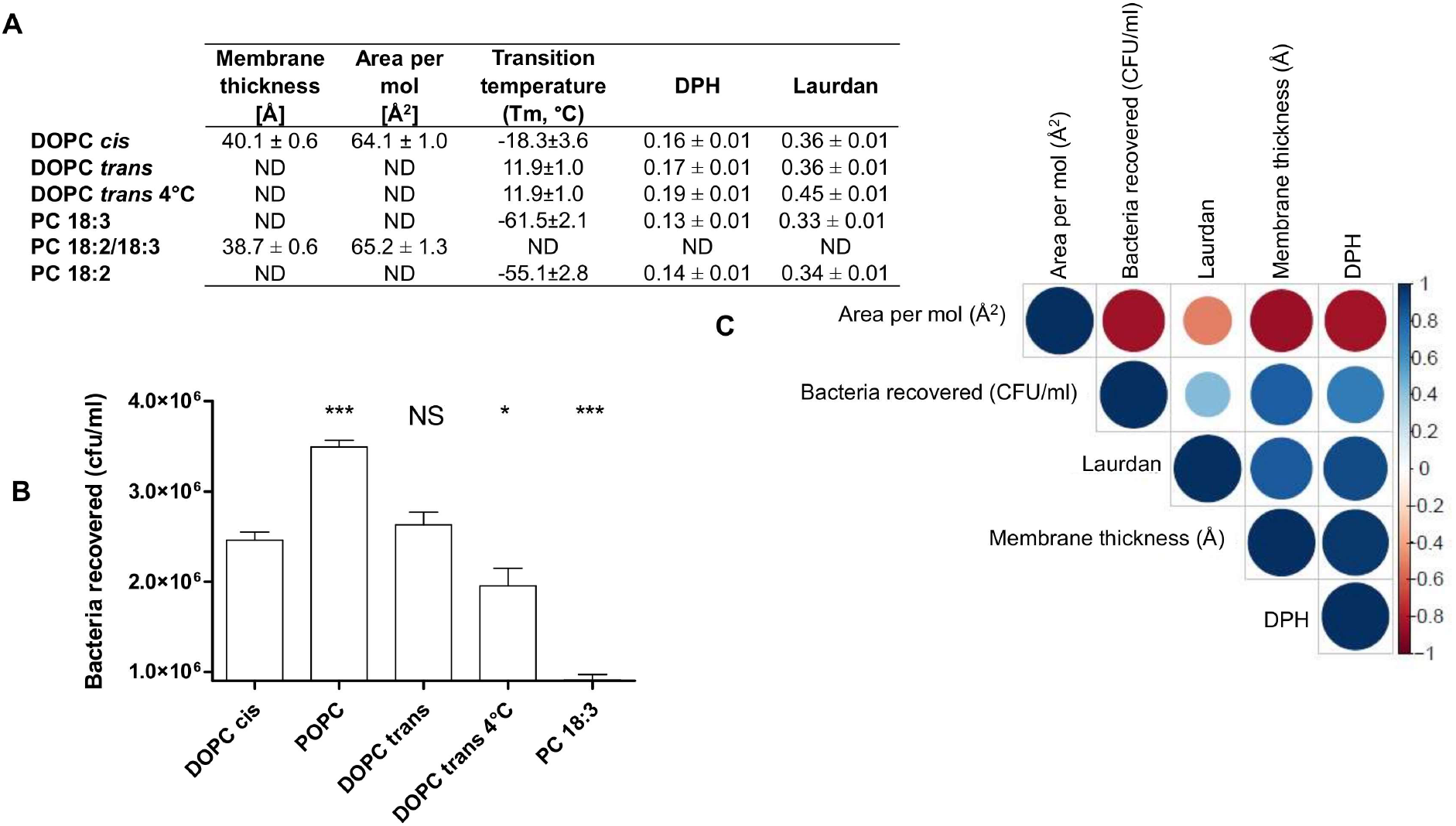
Impact of fatty acids with phosphatidylcholine (PC) head group on EHEC flagella adhesion. (A) EHEC adhesion on PC with different fatty acids composition. All lipids were tested pure. Non-specific adhesion on plastic was subtracted. DOPC cis: n=13; POPC: n=25; DOPC trans: n=13; DOPC trans 4°C: n=15; PC18:3: n=11 biologically independent samples. The bar graphs represent the mean of the reported data. Statistical significances were determined by a two-tailed Student’s *t* test. Significance levels are defined as follows: ***: *p* ≤ 0.001; *: *p* ≤ 0.05; NS: non-significant. (B) Table summarizing anisotropy values for diphenylhexatriene (DPH) and Laurdan probes at 23 °C and transition temperatures based on (32). The averages and standard deviations were calculated from at least three independent measurements. Membrane thickness and area per molecule, as well as related standard deviations, were calculated over the last 200 ns of MD simulations. (C) Correlogram representing Pearson’s correlation coefficient calculated for bacteria recovered, membrane thickness, area per mol, DHP and Laurdan anisotropies for the following lipids: DOPC cis, POPC, POPC:DOPC (from figure 4), PC18:3 and PC18:2 at 23°C. For PC18:3, membrane thickness and area per molecules are from PC18:2. The other parameters are from PC18:3. Positive correlations are visualized in blue while negative correlations are in red. The color intensity and the size are proportional to the correlation coefficients. On the right-hand side of the correlogram, the legend color shows the correlation coefficients and the corresponding colors. More detailed information on the significance of the correlations as well as on the correlation coefficients can be found in Fig. S7.

### Interactions of bacterial flagella with cell lines and lipid rafts

To provide further evidence about the impact of lipid composition with physiological membranes, colon epithelial cells (HT-29) enriched in saturated (palmitic acid) or unsaturated fatty acids (α-linolenic acid) were used. The lipid content of treated HT-29 cells was evaluated by GC-FID (Fig. 6A), which confirmed enrichment in both palmitic and α-linolenic acids. A LC-MS analysis confirmed these results and showed an accumulation of palmitic acid and linolenic acid when treated with the corresponding fatty acid (Fig. 6B). Palmitic acid was more incorporated into intracytoplasmic lipid droplets through triglycerides (TAG) than polyunsaturated fatty acid (C18:3) (33). Linolenic acid was found as well in molecular species associated with the plasma membrane (phospholipids). This result was described earlier with another human colon cell line (Caco-2 cells) (34). After the cells were washed, they were incubated with EHEC WT or a Δ*fliC* mutant for 30 min. As with biomimetic membranes, significantly less bacteria adhered on HT-29 treated with α-linolenic acid probably because of the effect on plasma membrane fluidity of this fatty acid. This result was observed only when EHEC were flagellated, highlighting the role of flagella in the adhesion process (Fig. 6C) with a significant impact of membrane fluidity on the strength of flagellum-membrane binding.

**FIG 6.**
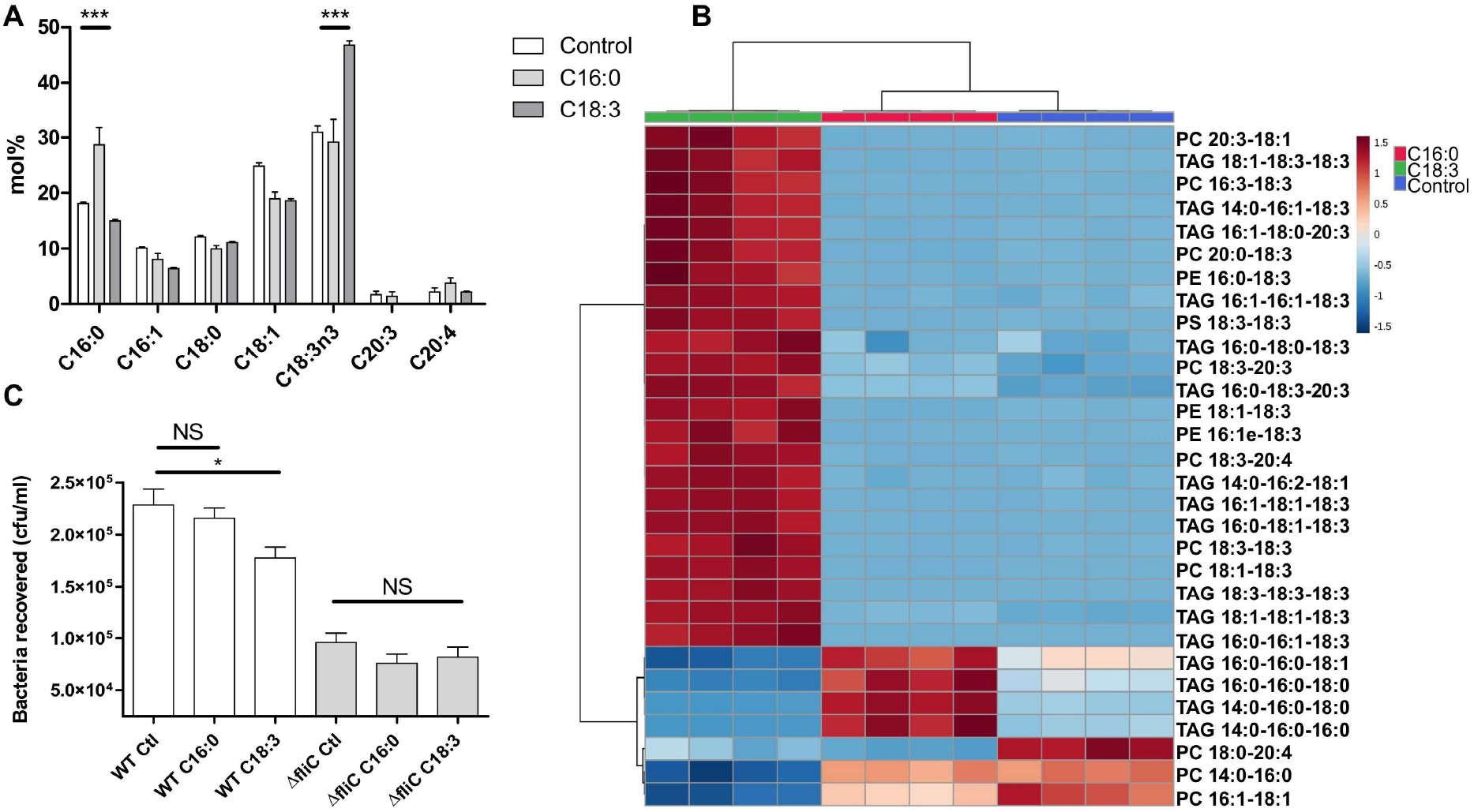
Modulation of fatty acid content in HT-29 and effect on EHEC adhesion. (A) Fatty acid methyl esters analysis of HT-29 cells by GC-FID, according to their diet: ethanol (control), C16:0 or C18:3. n=4 biologically independent samples. Statistical significances were determined by a two-tailed Student’s *t* test. (B) Heatmap of the 30 most statistically different lipids species (right) analyzed by LC-MS comparing the control (ethanol, blue) and the palmitic acid (C16:0, red) and the α-linolenic acid treated cells (C18:3, green). Colour coding indicates greater deviation from the mean of all samples for a particular lipid. The analysis was performed using MetaboAnalyst V4.0. (C) Adhesion of EHEC WT and Δ*fliC* on HT-29 after the different treatments. n=12 biologically independent samples. The bar graphs represent the mean of the reported data. Statistical significances were determined by a two-tailed Student’s *t* test. Significance levels are defined as follows: ***: *p* ≤ 0.001; *: *p* ≤ 0.05; NS: non-significant.

Based on these observations, we next addressed the role of plasma membrane fluidity with membrane models mimicking lipid rafts as they have been suggested to play a role in bacterial adhesion (19, 35) or in *E. coli* flagellar adhesion (20) on epithelial cells. Three lipids were used for these experiments: POPC, cholesterol (chol) and palmitoylsphingomyelin (PSM). This ternary lipid system is a model for lipid rafts with a phase diagram established at 23 °C (36). The ratios of the three lipids were set so that three different lipid orders were obtained, namely the liquid ordered (lo) (POPC/PSM/chol, 24.87:36.00:39.13 molar ratio), liquid disordered (ld) (POPC/PSM/chol, 71.53:23.25:5.22 molar ratio) and a mixed lo/ld system (POPC/PSM/chol, 1:1:1 molar ratio) (Fig. 7A). It is currently accepted that rafts are the cellular equivalent of lo phase *in vitro* (37). The correlations between the area per lipids, the membrane thickness, anisotropy and WT adhesion highlighted the most significant correlation between anisotropy and adhesion. As expected for lo, fewer adhering flagellated bacteria were observed in respect to POPC only, due to the gel phase state of the bilayer. The coexistence of lo and ld phases improved adhesion of the WT. In the ld phase, the ternary lipid system exhibited no significant differences with respect to POPC only. However, except for ld phase, non-flagellated bacteria were clearly less adherent. The fact that no significant difference was observed for ld when comparing the WT to the Δ*fliC* mutant, can be attributed to the presence of other adhesins capable of binding to chol or PSM (Fig. 7B). Correlations between WT adhesion and the parameters determined by MD and anisotropies, give the best result with the fluidity state with a moderate negative correlation (Fig. 7C and 7D). These results substantiate the previous hypothesis concerning an ideal plasma membrane fluidity to increase bacterial adhesion on lipid vesicles. However, with the ternary lipid system, the membrane is not homogeneous, which provides different sizes of lipid rafts depending on the lipid proportion. lo does not have rafts, lo/ld contains large rafts (>75-100nm) and ld forms small rafts (<20nm) (38).

**FIG 7.**
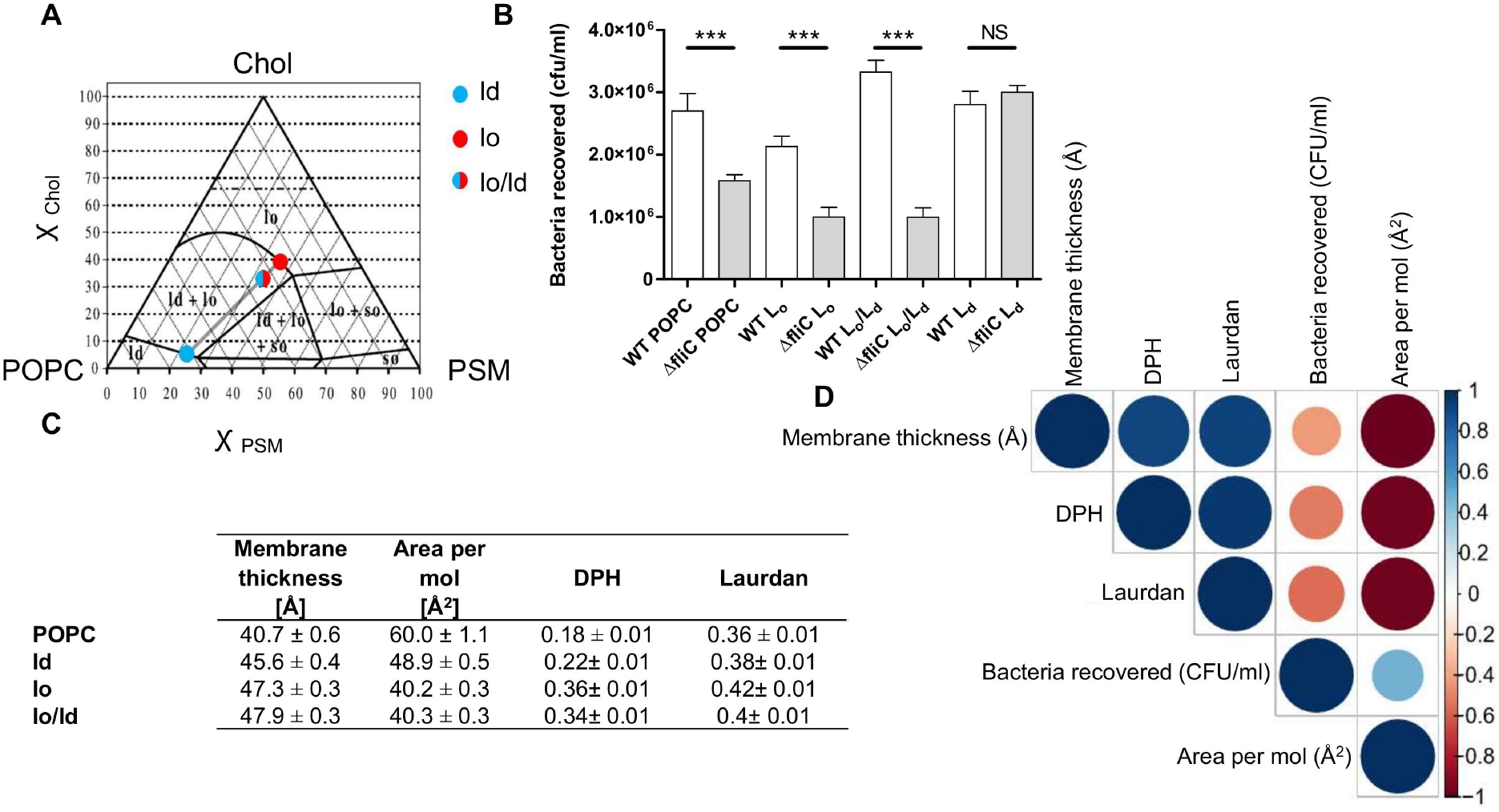
EHEC adhesion and lipid rafts. (A) POPC/PSM/chol phase diagram at 23 °C adapted from (36). The blue, red and blue/red circles are respectively ld, lo and lo/ld. The dashed horizontal line for χ_Chol_ 0.66 represents the cholesterol solubility limit on the lipid bilayer. (B) Adhesion of EHEC WT and Δ*fliC* on three different lipid mix from the ternary lipid system (POPC/PSM/chol). n=36 biologically independent samples for all Δ*fliC* and WT, except WT lo/ld n=30 biologically independent samples. The bar graphs represent the mean of the reported data. Statistical significances were determined by a two-tailed Student’s *t* test. Significance levels are defined as follows: ***: *p* ≤ 0.001; NS: non-significant. Nonspecific adhesion on plastic was subtracted. The average and standard deviation are calculated from three independent experiments. (C) Table summarizing anisotropy values for diphenylhexatriene (DPH) and Laurdan probes at 23 °C. The averages and standard deviations were calculated from at least three independent measurements. Membrane thickness and area per molecule, as well as related standard deviations, were calculated over the last 200 ns of MD simulations.(D) Correlogram representing Pearson’s correlation coefficient calculated for bacteria recovered, membrane thickness, area per mol, DHP and Laurdan anisotropies for the following lipids: POPC, ld, lo and lo/ld. Positive correlations are visualised in blue while negative correlations are in red. The color intensity and the size are proportional to the correlation coefficients. On the right-hand side of the correlogram, the legend color shows the correlation coefficients and the corresponding colors. More detailed information on the significance of the correlations as well as on the correlation coefficients can be found in Fig. S8.

## Discussion

Interactions with the plasma membrane are crucial for bacterial adhesion. Until recently, only a few studies have paid attention to enteric pathogens infection and plasma membrane lipid content during host invasion (35). Most of studies have focused on proteins as target for bacterial adhesion (21). By studying the flagellum-driven bacterial adhesion on biomimetic lipid bilayer membranes, we evidenced that the flagella of EHEC fully exploit the lipid bilayer to adhere. This interaction was originally identified to occur with negatively charged phospholipids by using immobilized lipids and purified flagella on a thin layer chromatography, which are not physiologically organized (15). Likewise, S. *enterica* serorvar Typhimurium *(Salmonella* Typhimurium) flagella were described to interact with pure cholesterol coated on surfaces but not organized in the complex structure of a plasma membrane (39). However, recently, we found that methylated flagella of *Salmonella* Typhimurium facilitate bacterial adhesion to PC GUV and negatively affect adhesion on pure POPG GUV (16).

The present work illustrates both the importance of flagellar motility, and the size and resulting curvature of the lipid vesicles, as important factors for optimal bacterial adhesion. Until now, the fatty acid composition was largely ignored but this work reveals its key role. Fatty acid saturation strongly impacts membrane thickness and area per lipids, both parameters thus appear as important biomarkers of flagellar adhesion. In the presence of saturated fatty acid, flagellar adhesion is optimal and modulated by the head group moiety of the membrane lipids. Conversely, the presence of unsaturated fatty acids decreases bacterial adhesion (Fig. 5), an effect which is even stronger when using polyunsaturated fatty acids (PC 18:3), where almost no flagellar adhesion was observed. The effect of polyunsaturated fatty acids on flagellar adhesion was still relevant on more complex plasma membranes, which includes proteins, such as a human cell line (Fig. 6). These results reflect a clear role of lipid packing on flagellar adhesion.

The computed bilayer parameters, obtained by MD simulations, such as membrane thickness and area per molecule, have been related to the membrane ordering and fluidity (40). A more fluid membrane exhibits less order and has higher value of area per lipid-molecule and lower value of membrane thickness (when comparing lipid chains of same size). A global correlation between these two structural parameters and the fluidity state of the GUV bilayer showed a very high level of correlation with most of the bilayer composition used in this study. In turn, these two parameters correlated with bacterial adhesion. Although not perfect because many other parameters are at stake, these two parameters provide a very good trend about the optimal conditions that increase, or not, bacterial adhesion. In other words, these two parameters, which can be computed at a relatively low computational cost, are easily obtained descriptors of the bacterial adhesion process. By adding a few other descriptors, we believe that we could establish a quantitative structure activity relationship (QSAR) that could predict flagellar adhesion with high robustness.

Ectothermic organisms incorporate fatty acids in phospholipids *via* a mechanism termed “homeoviscous adaptation” to have constant viscosities at the temperature of cell growth (41). It is only within homeoviscous adaptation limits that the diet of organisms can influence membrane lipid profile. In endothermic animals, the membrane composition has been thoroughly documented to be influenced by dietary fats in the erythrocyte plasma membrane and later in the liver, the brain and other organs (42–45). However, the influence of dietary intake on plasma membrane composition depends on the type of fatty acids ingested. Omega-3 and omega-6 fatty acids are not produced in mammals due to the lack of *ad hoc* desaturases, therefore humans must consume such fatty acids (46). In contrast, palmitic acid can be synthesized endogenously and its quantity is controlled under normal physiological conditions in order to not affect the membrane properties (47). It is noteworthy that the incorporation of diet fatty acids into membranes is influenced by the omega-3/omega-6 balance (48). To successfully invade hosts, EHEC and *Salmonella* Typhimurium require adhesion onto the intestinal epithelium (49, 50). Again, this process depends on plasma membrane fatty acid composition controlled by the diet (51). The impact of polyunsaturated fatty acids on human health during bacterial infection focuses on inflammation. α-linolenic acid is converted in arachidonic acid-derived inflammatory eicosanoids (52–54). Omega-3 fatty acids are converted as well to bioactive mediators, termed specialized pro-resolving mediators, that actively reprogram the host immune response to limit inflammation (55). However, little is known about the biophysical impact on bacterial adhesion and colonization of these fatty acids of the plasma membrane (35). Our results support the idea that a fatty acid diet can affect human health and, more surprisingly, host susceptibility to enteric pathogens by influencing flagellar adhesion on the plasma membrane.

Recently, bacterial flagellar motility was described to help bacteria to reach preferred sites in host plasma membrane containing sphingolipid-rich domains (56). Our findings show that not only motility but also bacterial flagella *per se,* acting as an adhesin on lipids, help bacteria to reach their host cell targets. As lipid rafts are the most documented plasma membrane lipids involved in bacterial adhesion (19, 22), we investigated three different types of lipid rafts: lo, lo+ld and ld, reflecting no, large and small size rafts respectively (38). No and small rafts did not substantially facilitate flagellar adhesion whereas large size rafts promoted adhesion (Fig. 7). These results suggest that an optimal size of lipid rafts exists that allows flagellar adhesion, which is consistent with previous works describing lipid rafts and bacterial adhesion (19, 22).

While our results highlight a previously unrecognized advantage of flagellated bacteria over bacteria lacking flagella (like *Shigella spp.* or *Staphylococcus spp.)* in host cell adhesion, the presence of flagellin can be deleterious by inducing the immune system (10). To avoid this major drawback flagellated bacteria down-regulate flagellin expression during planktonic/sessile switch and host invasion leading to immune evasion (57, 58). Accordingly, it is reasonable to speculate that, for at least EHEC O157:H7 and *Salmonella* Typhimurium (16), this risk is compensated by the capacity of the flagella to facilitate adhesion of the bacteria to the host cell plasma membrane. The increased adhesive properties of flagellated bacteria might increase the capability of the bacteria to invade their host. Furthermore, a recent paper illustrated the possibility that enzymatic or other unknown functionalities can be incorporated into the D3 domain of flagellins (59). Future work to explore lipolytic flagellins or other functional properties associated to lipids in flagellated bacterial species will be needed.

## Material and methods

### Bacteria growth and preparation

EHEC O157:H7 TUV93-0 derived from strain EDL933 was used (60).All The isogenic mutants Δ*fliC* and Δ*motA* were obtained as described (61). All strains were grown in lysogeny broth (LB) medium (1% w/v tryptone, 0.5% yeast extract and 0.5% NaCl) overnight at 37°C and 100 rpm. Bacterial motility was subsequently checked at 37°C after 7 h on motility medium (1% w/v tryptone, 0.33 w/v agar and 0.4% w/v NaCl). For all experiments, the overnight culture was centrifuged at 3500 *g* for 15 min at 20°C, resuspended in HEPES buffered saline solution (HEPES 20 mM pH 7.4, NaCl 150 mM) and diluted to a final concentration of 10^8^ UFC/ml.

### Liposome preparation

GUVs were prepared according to PVA-assisted gentle hydration (62). A 5% polyvinyl alcohol (PVA) solution (w/w) was prepared in water with 280 mM of sucrose and strongly stirred at 95°C until complete dissolution of PVA. 200 μl of PVA solution were spread manually onto a cover glass with a needle and dried on a heating plate at 50°C for 30 min. 5 μl of lipid (Avanti Polar Lipids) solution in chloroform at 3 mg/ml were subsequently deposited four times and spread until solvent evaporation. The residual solvent that could remain on the lipid-coated cover glass is evaporated under vacuum at least 1 h. To form a well on the cover glass slide, a ring was glued onto it and 500 μl of HEPES buffered saline solution was added for hydration. After swelling of 1h at room temperature, the giant unilamellar vesicles (GUVs) formed were either directly stored in a fridge or reengage in hydration of another preparation to obtain a higher lipid concentration. Most of the GUVs size were evaluated manually under microscope (Fig. S1).

Another method of formation can be applied in order to form vesicles with smaller diameter: from Large Unilamellar Vesicles (LUV) to small GUVs. The lipid solution in chloroform was dried under a nitrogen stream, and then under vacuum for 2 h to remove remaining solvent. The film was hydrated to the desired lipid concentration in HEPES buffered saline solution. After vortexing, the multilamellar vesicle suspension was extruded 25 times using a syringe type extruder with polycarbonate filter having a pore size of 400 nm for 400-nm liposome, 2 μm for 1 μm-liposome and 5 μm for 2 μm-liposome (Liposofast, Avestin Inc). Prior extrusion for 400-nm liposome, the solution was sonicated using a tip sonicator (Ultrasonic Processor Vibra Cell, Sonic Materials). Liposome size was determined either manually with ImageJ by epifluorescence microscopy or by dynamic light scattering (Zetasizer Nano ZS, Malvern Instruments) for liposomes with a diameter smaller than 1 μm.

### Labelling and epifluorescence microscopy

Liposomes were first immobilized on gold-coating glass. Prior to liposome preparation, DSPE-PEG-PDP (Avanti Polar Lipids) was added to the lipid mixture in chloroform at 3%w and was as well doped with 2% mol NBD-PE for liposome labelling (Avanti Polar Lipids). At the same time, a cover slip was coated with 1 nm of chromium and 10 nm of gold by thermal evaporation (Evaporator Edwards model Auto 306). The incubation between liposomes and bacteria was done in a separated container for 1 h and transferred onto a microscope chamber. This latter was composed of the gold-coated glass surface at the bottom, spaced from a common microscope slide with lateral spacers of molten Parafilm. Observation was carried out on Leica DMI6000 B epifluorescence microscope.

### Bacterial adhesion assays on liposomes

Bacterial adhesion assays were done in 6-well plates (9.6 cm^2^ per well). Gold-coated slides of 1.9×2.5 cm were placed onto a 3 mm high pedestal in each well. 1 ml of 5 μg/ml liposome solution containing DSPE-PEG-PDP was homogeneously deposited on gold-coated surface and incubated for 1 h. After adding 10 ml of HEPES buffered saline solution, the pedestal was carefully removed with a bended pipette tip. A volume of 8 ml was then removed to eliminate non-immobilized liposomes with a minimal volume, to keep immerged the gold-coated surface. The immobilized liposomes were incubated for 1 h with 5 ml of bacterial suspension at 10^8^ UFC/ml in HEPES buffered saline. Prior to discard all medium, the excess of bacteria was removed by taking away 5 ml and the non-adherent bacteria were removed from the wells with 8 ml of fresh HEPES buffered saline. With 1 ml of PBS, adherent bacteria were detached by pipetting vigorously several times directly onto gold-coated surface. Samples were serially diluted and plated on LB agar for viable bacterial counts. The bacterial count for 0 μg of lipid corresponds to the nonspecific bacterial adhesion onto the gold surface without lipids. The non-specific adhesion of a well alone (without glass and lipids) from the 6-well cell-culture plate use to perform the assay was systematically subtracted. All adherence assays were performed at 23 ± 2 °C and at 4°C for DOPC (Δ9-trans).

### Bacterial adhesion assays on HT-29

The human colonic cell line HT-29 was obtained from the American Tissue Culture Collection (ATCC). The cell line was maintained in modified McCoy medium supplemented with 10% (v/v) heat-in-activated fetal calf serum, 2mM L-glutamine, 100 unit/ml penicillin and 100 unit/ml streptomycin at 37 °C in 5% CO2. Cultures were used between passages 15 to 20. The cells were seeded in 24-well culture plates (1.9 cm^2^ per well) at a concentration of 4 × 10^4^ cells per well. The culture medium was changed every day. After 24 hours, the cells were treated with 100 μM of C18:3 (α-linolenic), C16:0 solubilized in ethanol or only ethanol as control. After 24 hours, the supernatant was removed and 2 × 107 bacteria in 250 μl in PBS were incubated for 30 min at 37°C. Then the cells were rinsed twice with PBS and treated with trypsin-EDTA. After samples dilution, samples were plated on LB agar for viable bacterial counts.

### Laurdan generalized polarization measurements

1 mM Laurdan (Sigma-Aldrich) solution in DMSO was added to vesicle suspension, in order to have a lipid:probe ratio of 20:1. This mixture was incubated in the dark for 30 min and further diluted in HEPES buffered saline to a final concentration of 5 μM of Laurdan and 100 μM of phospholipids. Flagella were purified as previously described by mechanical shearing (15). Increasing quantities of H7 flagella were progressively added to the Laurdan/liposomes mixture and incubated each time in the dark for 20 min in a circulating water bath at 37°C. Generalized Polarization (GP) (23) was calculated from the emission intensities at 440 nm and 490 nm after an excitation at 390 nm, according to the following equation (I):

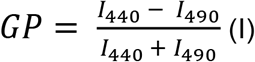

using Varian Cary Eclipse fluorescence spectrophotometer (Agilent Technologies). The relative ΔGP is obtained by subtracting the GP value in the absence of flagella from all GP values, for each size of liposome.

### Fluorescence anisotropy measurements

Steady-state fluorescence emission anisotropy (r) was measured with a Cary Eclipse Fluorescence Spectrophotometer (Agilent Technologies) equipped with a thermostated cuvette holder. GUVs with an average diameter of 2 μm and made of different lipid mixtures were prepared from a lipid dried film by extrusion as previously described. The fluorescent probes 1,6-diphenyl-1,3,5-hexatriene (DPH) or Laurdan fluorescent probes (Sigma-Aldrich) were added prior the lipid film formation at the lipid:probe molar ratio of 40:1. The GUVs were diluted in HBS solution to the final concentration of 200 μM of lipids and 5 μM of fluorescent probes in the quartz cuvette and were incubated at the desired temperature for 30 min in the dark prior to fluorescence measurement. Fluorescence intensities were collected at 435 nm for DPH and for Laurdan with excitation wavelengths 357 and 360 nm respectively. Anisotropy was automatically calculated by the software of the spectrophotometer according to Equation (II):

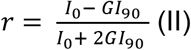

where *I*_0_ is the fluorescence intensity measured with polarizer in parallel orientation (0°) and *I*_90_ the intensity in perpendicular orientation (excitation 0° and emission 90°). *G* is the correction factor derived from the ratio of emission intensity at 0 and 90° with the excitation polarizer at 90° and is taking into account the different sensitivity of the detection system for vertically and horizontally polarized light (Equation III):

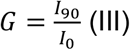

### NMR for lipid ratio calculation in GUVs

High resolution ^1^H spectra were recorded on a Bruker AVIII NMR spectrometer equipped with a 5 mm BBFO probe head operating at 400 MHz ^1^H Larmor frequency. All experiments were carried out at room temperature. The instrument’s standard pulse sequence program (zg30) was used with the following parameters: 2s acquisition time, 20 ppm spectral width, 5 s relaxation delay. Deuterated chloroform and 5 mm NMR tubes were purchased from Eurisotop (Saint-Aubin, France). In total, 3072 repetition experiments were performed leading to a total acquisition time of 6 hours. After data processing, phase and baseline correction, the area of the peaks of interest was determined by integration and the molar ratio of lipids was calculated with relative integrations.

### GC-FID

Fatty acids from seeds, seedlings and plant tissues were directly transmethylated as described before (63). Heptadecanoic acid (C17:0) was used as an internal standard. Fatty acids methyl esters were extracted in heptane and separated by gas chromatography with flame ionization detection.

### Lipids extraction

The fatty acid content of the cells or GUVs was determined after extraction with a modified method according to Bligh and Dyer (64). 375 μl of a mixture of chloroform/methanol (2:1, v:v) containing BHT (1 mM) was added to 100 μl of aqueous solution containing lipid vesicles or cells. The mix was vortexed during 10-15 min, then 125 μl of chloroform was added and mixed 1 min. Followed by 125 μl of water and centrifuged quickly. The lower phase was collected, dried under a stream of nitrogen at room temperature and kept at −20°C until needed. Prior liquid chromatography/high resolution tandem mass spectrometry (LC-HRMS^2^) analyses

### LC-HRMS^2^ analyses

The lipids were resuspended in 200 μl of isopropanol. LC was performed based on a modified protocol from (65). Briefly an HPLC 1290 (Agilent Technologies) with a C18 Hypersil Gold (100 x 2.1 mm, 1.9 μm, Thermofisher) at 50°C was used. Mobile phases were 60:40 (v/v) acetonitrile:water with 10 mM of ammonium formate and 0.1% formic acid (solvent A) and 90:8:2 (v/v) isopropanol:acetonitrile:water with 10 mM of ammonium formate and 0.1% formic acid (solvent B). The gradient was as follows: A = 0–2 min 68%, 2–8 min 60%, 8–10 min 55%, 10–16 min 50%, 16–22 min 40%, 22–28 min 30%, 28–35 min 20%, 35–40 min 0%. Finally, the column was equilibrated for 6 min with 68%. The flow rate was set at 0.25 ml/min and with an injection volume of 2 μl. LC-ESI-HRMS2 analyzes were achieved by coupling the LC system to a hybrid quadrupole time-of-flight (QToF) mass spectrometer Agilent 6538 (Agilent Technologies) equipped with a Dual electrospray ionization (ESI). The source temperature, fragmentor and the skimmer were set up respectively at 350°C, 150 V, 65 V. The acquisition was made in full scan mode between 100 m/z and 1700 m/z, with a scan of two spectra per second. Selected parent ions were fragmented with collision energy fixed at 35 eV. MS2 scans was done on the sixth most intense ions. Two internal reference masses were used for in-run calibration of the mass spectrometer (121.0509, 922.0098 in positive ion mode and 112.9856, 1033.9881 in negative ion mode). MassHunter B.07 software allowed to control the parameters of the machine, acquired and processed the data. The mass spectra were acquired in positive and negative-ion mode.

#### Data processing and annotation

Agilent generated files (*.d) were converted to *.mzXML format using MSConvert (66). Files *.mzXML datasets were processed using MZmine 2 v2.37 (67). The noise level was 2.0E3 for MS1 and 0E00 for MS2 in centroid. The chromatogram builder was used using a minimum time span of 0.10 (min), a minimum of height of 1.0E3 and m/z tolerance of 5ppm. The chromatogram deconvolution was done with the local minimum search algorithm. The chromatographic threshold was 30.0%; the search minimum in RT range was 0.05 min with a minimum relative height of 5% and a minimum ratio of peak top/edge of 2. Peak duration range 0.05 – 3 min. MS2 scans were paired using a m/z tolerance range of 0.05 Da and RT tolerance range of 0.1 min. Isotopologues were grouped using the isotopic peaks grouper algorithm with a m/z tolerance of 0.008 and a RT tolerance of 0.3 min. A peak alignment step was performed using the join aligner module (m/z tolerance = 0.008, weight for m/z = 50, weight for RT = 50, absolute RT tolerance 2 min). Peak finder module was used with intensity tolerance of 10%, m/z tolerance of 0.008 and retention time tolerance of 1.0 min. The resulting peak list was then filtered using the peak list row filter module with a minimum peak in a row of 2, a minimum peak in an isotope pattern of 2 and by keeping only peaks with MS2 scan (GNPS). The peak list was then exported to *.csv using the module “Export to CSV file”. Moreover, a *mgf was exported using the module: “export for/submit to GNPS”. The peak list was annotated using a combination of four databases, GNPS (68) lipid blast (69), lipid match (70) and lipiDex (71). Non-annotated features were removed.

### MD simulations

POPC and DOPC lipid bilayers were constituted of 128 lipid molecules in total and were put into a hydrated simulation box, as generated by CHARMM-GUI membrane builder (72). The mixtures (POPC/POPE, POPC/POPG, POPC/DOPC, POPC/DOPE and POPC/DOPG) were prepared accordingly, except that a 38:26 ratio was used in each leaflet to create the desired 40:60 molar ratio. To simulate the properties of the PC 18:3 membrane, the PC C18:2/C18:3 lipid was selected as being structurally close in terms of lipid tails and similar in the head group, and because this lipid has already been parameterised in the Charmm force-field, whereas PC 18:3 has not been yet. In the simulations with sphingomyelin, the ratios of POPC/PSM/CHOL were 21:22:21 for the simulation with equal ratios, 16:23:25 for the ordered phase and 43:14:4 for the disordered phase. 150 mM of NaCl plus ions necessary for system charge neutralisation were added to the simulation box to model the ion physiological concentration. The CHARMM-GUI charmm force field topology and parameter files were transferred to the amber format using parmed. The systems were minimised using the steepest gradient for 500 steps and conjugate gradient for 500 steps with the lipid molecules restrained, followed by minimising and heating. The production simulation was run for 500 ns at 293.15 K, using the Amber18 software package (73), with the Charmm36 forcefield for lipids and the TIP3P model for water (74), and with a time step of 2 fs. The SHAKE algorithm was used for hydrogen atoms (75). The Langevin thermostat and constant pressure periodic boundary conditions with anisotropic pressure coupling in the xy direction were applied. The last 200ns of each simulation was used to calculate all parameters (area per lipid, membrane thickness, and order parameters) as well as related standard deviations.

### Statistical analysis

For bacterial adhesion assays, the means ± S.E. were calculated from values obtained from five replicates and performed with at least three independent biological experiments. Three assays were carried out for lipid analysis. Statistical evaluation was performed with GraphPad Prism (GraphPad Software, San Diego California, www.graphpad.com). The statistical significance was evaluated with Student’s *t* test or oneway analysis of variance. The heatmap was performed on Metaboanalyst (76).The results were considered as significant for a *p* value≤0.05. Secondary data analysis and correlation matrices were calculated using the open-source software GNU-R. Bacterial growth counts across all the tested scenarios as well as metrics related to each lipid composition (viz. Membrane thickness, Area per molecule, DPH and Laudan anisotropy) were loaded as csv files into the R environment. Correlations were computed using the “`corr`” function of the R library “`Hmisc`”, which calculates the significance level (p-value) using Pearson correlation coefficient on raw values. The correlation matrix plots were generated using the R packages “`corrpiot`” and “`PerformanceAnaiytics`”. To interpret the size of the correlation coefficient it was as follow: 1.00-0.9, very high; 0.9-0.7, high; 0.7-0.5, moderate; 0.5-0.3, low; 0.3-0, negligible correlation as described (77).

## Acknowledgement

We are grateful to David Gally and Eliza Wolfson for Stx-negative derivative strain of TUV 93-0 and its isogenic mutants *(ΔfliC* and ΔmotA). We thank the European Regional Development fund ERDF and the Region of Picardy (CPER 2007-2020). H.C. and L.L. acknowledge support by the French Ministry of Higher Education, Research and Innovation. The authors would like to acknowledge Ashleigh Holmes for her critical review.

