## Supplemental fig 1 for "The impact of plasma membrane lipid composition on flagella-mediated adhesion of enterohemorrhagic *Escherichia coli*"

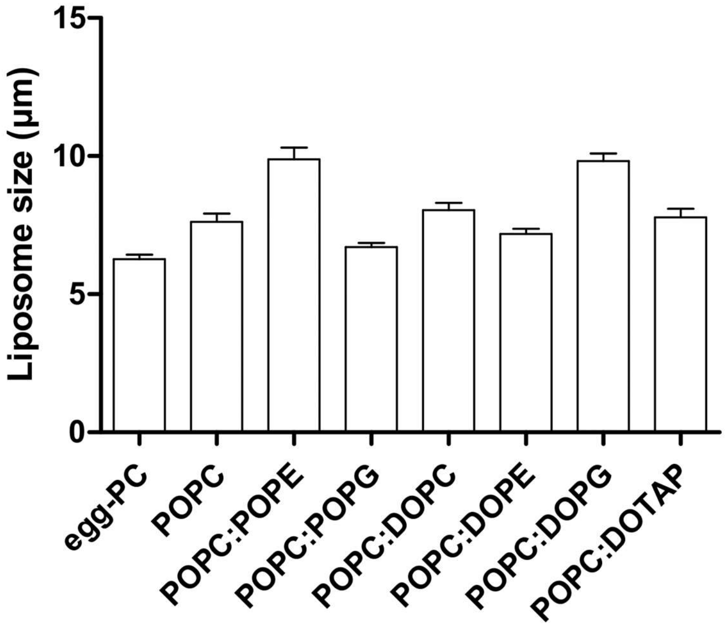


FIG S1

Sum-up of the liposomes sizes measured for the Fig. 4 and 5. Egg-PC: n=490; POPC: n=365; POPC:POPE: n=176; POPC:POPG: n=414; POPC:DOPC: n=333; POPC:DOPE: n=452; POPC:DOPG: n=339; POPC:DOTAP: n=283 GUVs measured. The bar graphs represent the mean of the reported data.
