## Supplemental fig 1 for "The impact of plasma membrane lipid composition on flagella-mediated adhesion of enterohemorrhagic *Escherichia coli*"

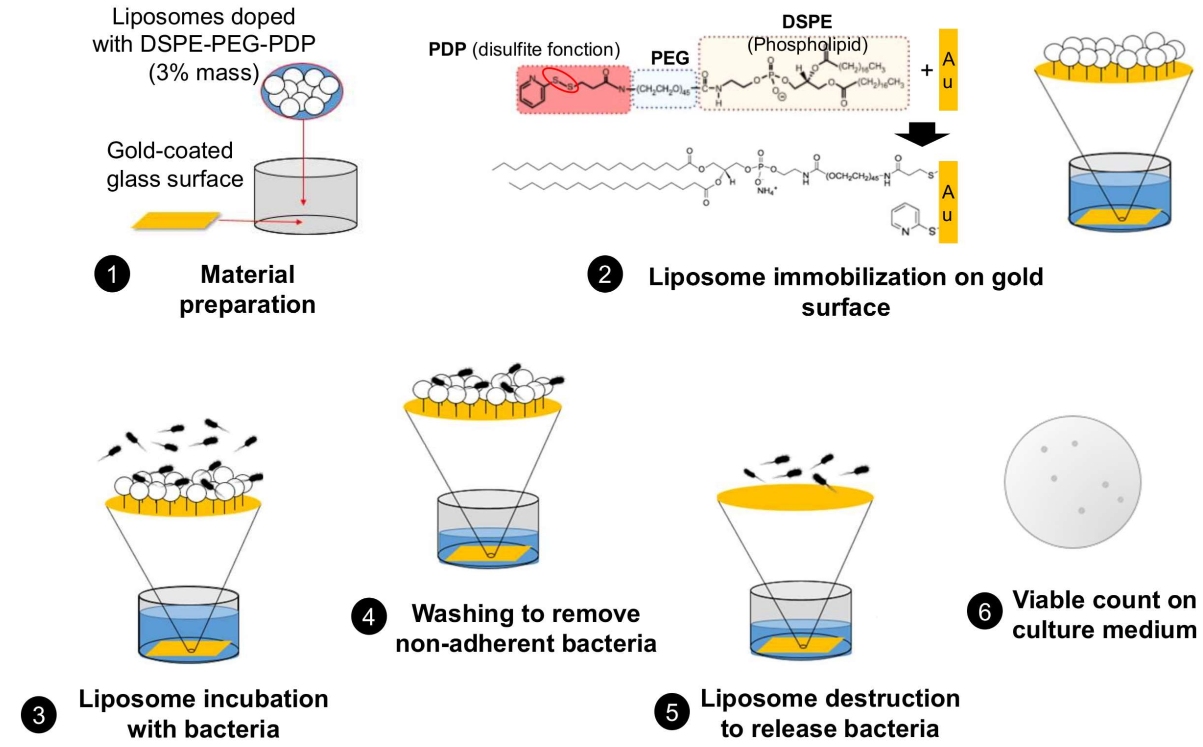


(DSPE-PEG-PDP): 1,2-distearoyl-sn-glycero-3-phosphoethanolamine-N-[pyridyldithiol propionate (polyethylene glycol)-2000]

FIG S2

Protocol for GUVs adhesion assay. (1) GUV and gold surfaces were prepared as described in the material and methods section. (2) 5 µg liposomes doped with DSPE-PEG-PDP (3%w) were immobilized on gold-coating glass surface in 6-well plates. (3) Liposomes were incubated during one hour with 5 ml of bacterial suspension at $10^8$ UFC/ml in HEPES buffered saline. (4) Non-adherent bacteria were removed from the wells with HEPES buffered saline. (5) Adherent bacteria were detached by pipetting vigorously several times with PBS directly onto gold-coated surface. (6) After serial dilutions, plating and 16hours of growth at 37°C, the bacteria were counted.
