## Supplemental fig 3 for "The impact of plasma membrane lipid composition on flagella-mediated adhesion of enterohemorrhagic *Escherichia coli*"

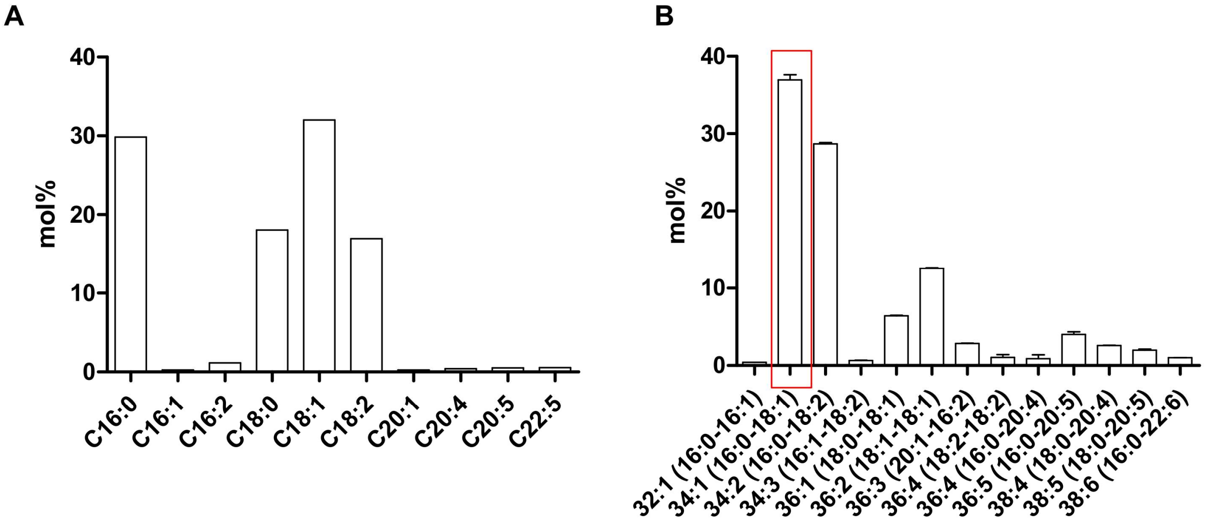


FIG S3

Egg-PC characterization. (A) Fatty acid methyl esters analysis of egg-PC by GC-FID. (B) PC species detected by LC-MS analysis of egg-PC used in our experiments. The most abundant lipid is encased in red (POPC). The bar graphs represent the mean of the reported data.
