## Supplemental fig 4 for "The impact of plasma membrane lipid composition on flagella-mediated adhesion of enterohemorrhagic *Escherichia coli*"

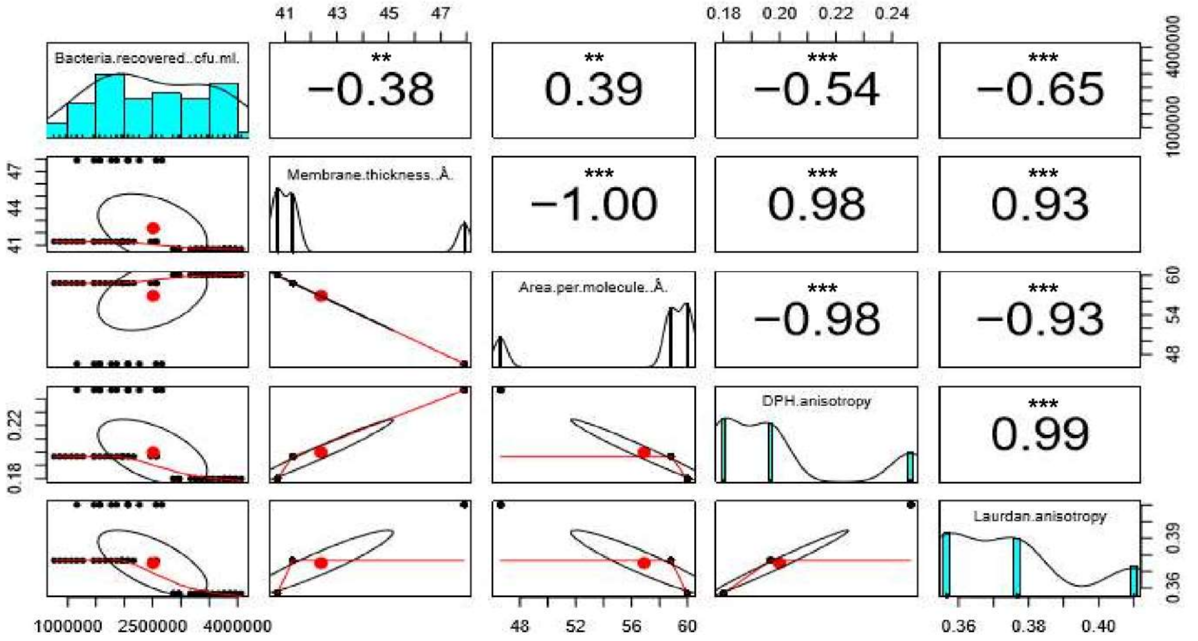


FIG S4

Corresponding to the Fig. 3: Impact of head polar phospholipids with one palmitic and one oleic acid (PO) on EHEC flagella adhesion. Corresponding scatter plots with the correlation coefficient (r) values and their significance. The distribution of each variable is shown on the diagonal subplots. A set of the bivariate scatter plots with a fitted line (displayed in red) are shown at the bottom of the diagonal Significance levels are defined as follows: ***: $p \leq 0.001$; **: $p \leq 0.005$.
