## Supplemental fig 6 for "The impact of plasma membrane lipid composition on flagella-mediated adhesion of enterohemorrhagic *Escherichia coli*"

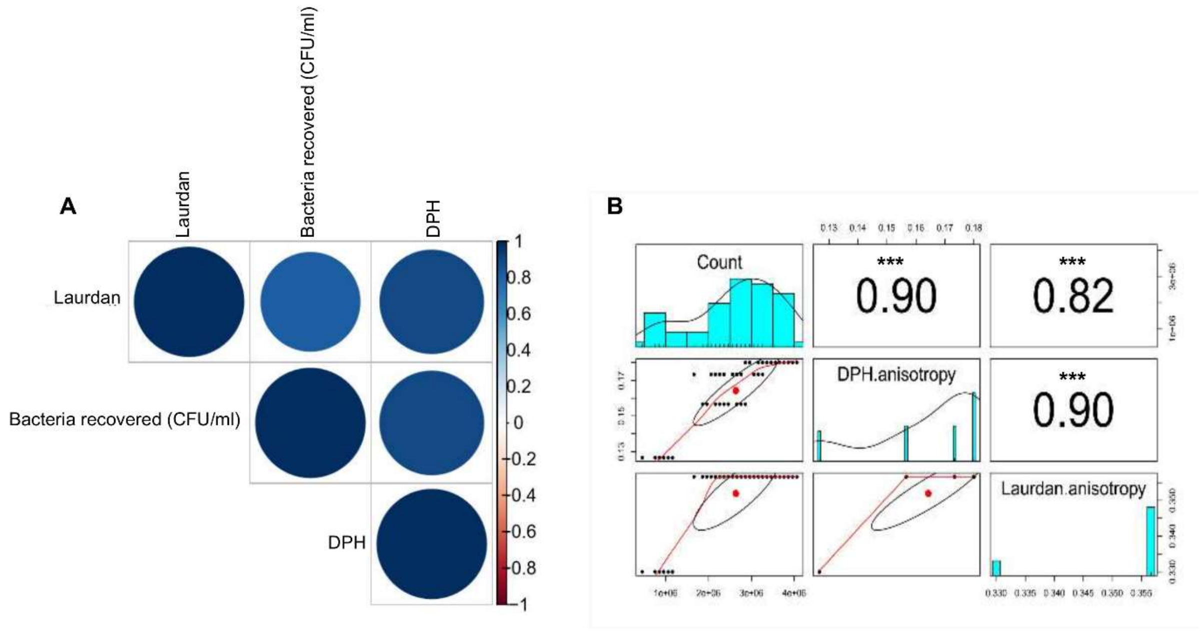


FIG S6

Fluidity and PC. Correlogram representing Pearson's correlation coefficient calculated for bacteria recovered, DHP and Laurdan anisotropies for the following lipids: POPC, DOPC cis DOPC trans and PC18:3 at 23°C. Positive correlations are visualised in blue while negative correlations are in red colour. The colour intensity and the size are proportional to the correlation coefficients. In the right side of the correlogram, the legend colour shows the correlation coefficients and the corresponding colours. (B) Corresponding scatter plots with the correlation coefficient (r) values and their significance. The distribution of each variable is shown on the diagonal subplots. A set of the bivariate scatter plots with a fitted line (displayed in red) are shown at the bottom of the diagonal Significance levels are defined as follows: ***: $p \leq 0.001$.
