## Supplemental fig 8 for "The impact of plasma membrane lipid composition on flagella-mediated adhesion of enterohemorrhagic *Escherichia coli*"

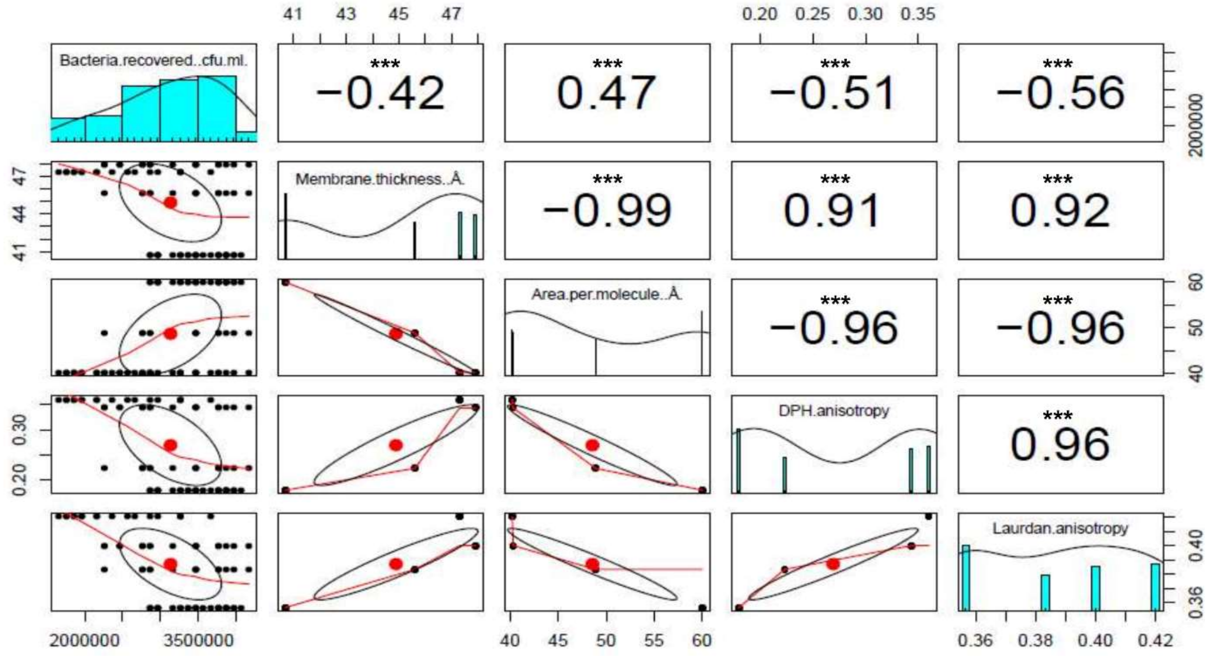


FIG S8

Corresponding to the Fig. 7: EHEC adhesion and lipid rafts. Corresponding scatter plots with the correlation coefficient (r) values and their significance. The distribution of each variable is shown on the diagonal subplots. A set of the bivariate scatter plots with a fitted line (displayed in red) are shown at the bottom of the diagonal Significance levels are defined as follows: ***: $p \leq 0.001$.
